# Protein cost minimization promotes the emergence of coenzyme redundancy

**DOI:** 10.1101/2021.05.17.444566

**Authors:** Joshua E. Goldford, Ashish B. George, Avi I. Flamholz, Daniel Segrè

## Abstract

Coenzymes distribute a variety of chemical moieties throughout cellular metabolism, participating in group (e.g., phosphate, acyl) and electron transfer. For a variety of reactions requiring acceptors or donors of specific resources, there often exist degenerate sets of molecules (e.g., NAD(H) and NADP(H)) that carry out similar functions. Although the physiological roles of various coenzyme systems are well established, it is unclear what selective pressures may have driven the emergence of coenzyme redundancy. Here we use genome-wide metabolic modeling approaches to decompose the selective pressures driving enzymatic specificity for either NAD(H) or NADP(H) in the metabolic network of *Escherichia coli*. We found that few enzymes are thermodynamically constrained to using a single coenzyme, and in principle, a metabolic network relying on only NAD(H) is feasible. However, structural and sequence analyses revealed widespread conservation of residues that retain selectivity for either NAD(H) or NADP(H), suggesting that additional forces may shape specificity. Using a model accounting for the cost of oxidoreductase enzyme expression, we found that coenzyme redundancy universally reduces the minimal amount of protein required to catalyze coenzyme-coupled reactions, inducing individual reactions to strongly prefer one coenzyme over another when reactions are near thermodynamic equilibrium. We propose that protein minimization generically promotes coenzyme redundancy, and that coenzymes typically thought to exist in a single pool (e.g., CoA) may exist in more than one form (e.g., dephospho-CoA).

**Significance statement:** Metabolism relies on a small class of molecules (coenzymes) that serve as universal donors and acceptors of key chemical groups and electrons. Although metabolic networks crucially depend on structurally redundant coenzymes (e.g., NAD(H) and NADP(H)) associated with different enzymes, the criteria that led to the emergence of this redundancy remain poorly understood. Our combination of modeling, and structural and sequence analysis indicates that coenzyme redundancy is not essential for metabolism, but rather an evolved strategy promoting efficient usage of enzymes when biochemical reactions are near equilibrium. Our work suggests that early metabolism may have operated with fewer coenzymes, and that adaptation for metabolic efficiency may have driven the rise of coenzyme diversity in living systems.

## Introduction

Group and electron transfer reactions form the foundation for more complex metabolic pathways in biological systems. For specific types of biochemical transformations, molecular components are shuttled within the cell through the action of small molecule coenzymes. For the majority of these transformations, there often exist multiple chemically distinct coenzymes capable of similar chemistry (Fig. 1a). For example, electron transfer between common functional groups widely distributed throughout cellular metabolism (e.g. alcohols, aldehydes, and activated carboxylic acids) are carried out by a pair of redundant coenzymes capable of electron transfer at identical standard thermodynamic potential: nicotinamide adenine dinucleotide (NAD^+^) and the phosphorylated derivative NADP^+^ (or the reduced forms, NADH and NADPH) (Fig. 1b). NAD(H) and NADP(H) are coupled to distinct sets of reactions, where NAD(H) is typically used in the breakdown of nutrients to generate precursors and ATP (catabolism), while NADP(H) is used in the reductive synthesis of macromolecules (anabolism) (1, 2). These functional roles have been attributed to differences in the *in vivo* thermodynamic potentials of NAD(H) and NADP(H), where NAD(H) and NADP(H) are typically poised in the oxidized (NAD^+^) and reduced state (NADPH), respectively (3–7). For example, in *E. coli* grown aerobically with glucose as the sole source of carbon, the NADH/NAD^+^ ratio is approximately 0.03, while the NADPH/NADP^+^ ratio exceeds 57 (3, 8). This thermodynamic difference is commonly assumed to be a universal way for cells to simultaneously oxidize and reduce metabolites that might be impossible with just a single nicotinamide coenzyme (2, 9).

**Figure 1:**
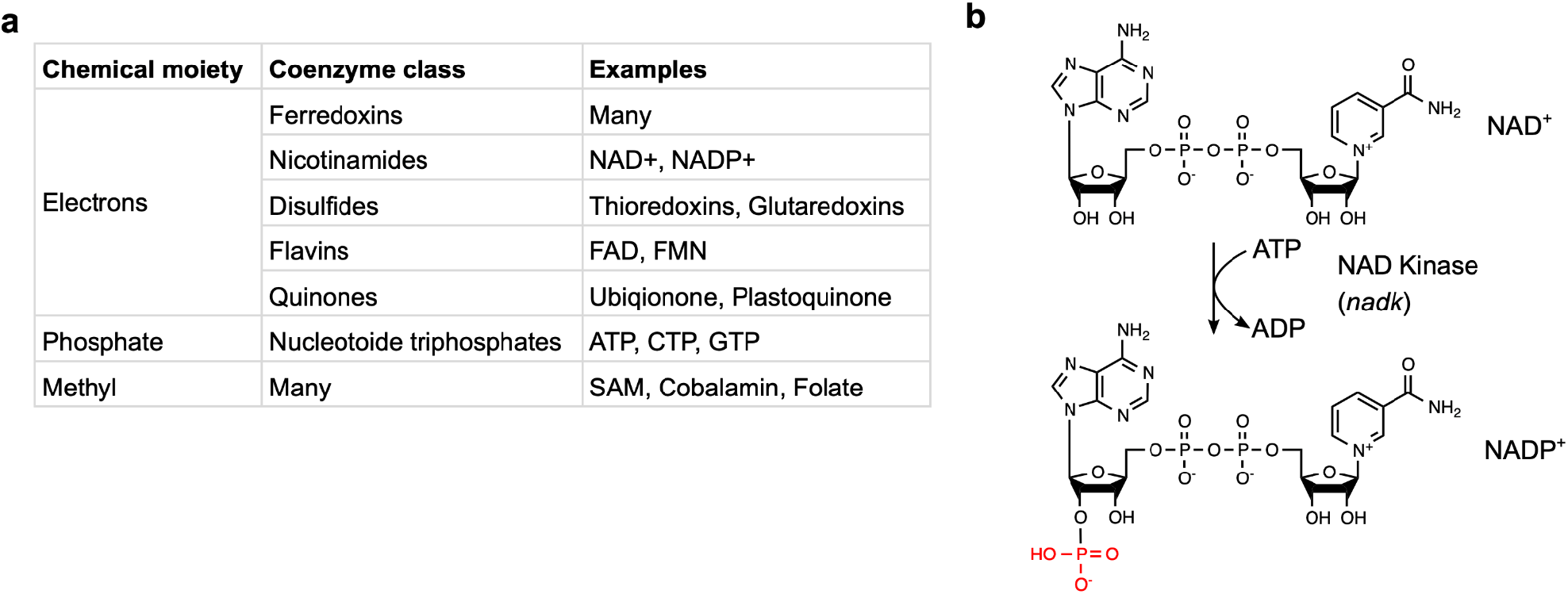
Coenzyme redundancy is a widespread biological phenomenon. **a**. A table of examples of redundant coenzymes for various classes of coenzymes. Note that for electron transfer reactions, different classes of coenzymes have varying characteristic ranges of standard redox potentials, enabling electron transfer between distinct classes of functional groups (27). **b**. Structure of the nicotinamide adenine diphosphate (NAD^+^) and the phosphorylated derivative (NADP^+^). The phosphoryl transfer enzyme NAD Kinase phosphorylates NAD^+^ on the 3’ hydroxyl to make NADP^+^.

The assumption that coenzyme redundancy is essential in living systems leaves unanswered a set of fundamental questions about the nature and early evolution of metabolism: is the coupling of each reaction to a specific coenzyme a strong constraint for the proper functioning of metabolism, or can some coenzymes be switched with limited consequences? More generally, is the presence of two coenzymes poised in different redox states an absolute necessity for the proper functioning of a metabolic network, or could metabolism in principle operate with only one coenzyme? If one coenzyme is sufficient, is it possible to quantitatively explain how evolution could gradually lead to the selective advantage of two coenzymes? And is the rise of multiple coenzymes an idiosyncratic step along a complex historical process, or a predictable universal feature of enzyme-driven biochemical networks?

While prior experimental work has shown that a range of cellular phenotypes can be sensitive to the coenzyme preferences for various oxidoreductases, they do not rule out the possibility of a simpler precursor to the NAD(H)/NADP(H) system. NAD(P)-dependent oxidoreductases can display selectivity for either NAD(H) or NADP(H), and perturbations to this specificity can strongly affect cellular phenotypes and fitness (10, 11). For example, mutagenesis experiments have shown that altering coenzyme specificity of isopropylmalate dehydrogenase and isocitrate dehydrogenase can severely reduce growth rate in *E. coli* (11–15). More generally, specificity for NAD(H) or NADP(H) for some enzymes can strongly influence intracellular reaction flux distributions (10), serving as the basis for metabolic engineering strategies to dramatically improve yields of valuable metabolic byproducts (10, 16–18). Importantly, recent work has demonstrated that specificity for NAD(P)H can emerge in laboratory evolution studies, suggesting that coenzyme preference might be a highly adaptable feature of metabolic networks (19). However, it is unclear if most enzymes are typically constrained to use only NAD(H) or NADP(H), and whether a single coenzyme could support the functioning of metabolism.

While an experimental assessment of the landscape of possible variants of metabolism with different coenzyme systems would be extremely challenging, one can leverage theoretical and computational approaches to help address these questions. The recent development and application of metabolic modeling has brought a quantitative and model-driven approach to the study of metabolic evolution (20–25), and has led to uncovering optimality criteria that may have shaped universal features of core metabolism, such as the emergence of both NAD(H) and NADP(H) (26). Although recent quantitative work has explored the logic for why NAD(P)H emerged as a prominent coenzyme in biochemistry(27), a quantitative theory predicting the emergence of both NAD(H) and NADP(H), rather than just a single coenzyme, remain unexplored.

Here we use computational systems biology and various network-level models of metabolism to systematically probe, in a way that could not be easily addressed experimentally, the possible advantages of maintaining multiple pools of biochemically redundant redox coenzymes. By taking advantage of large-scale computational experiments using genome-scale stoichiometric models, we show that the *Escherichia coli (E. coli)* metabolic network is predicted to be insensitive to coenzyme preference for the majority of enzymes, and that a metabolism with just a single coenzyme (rather than both NAD(H) and NADP(H)) is capable of producing biomass under a variety of environments. We found that switching coenzyme preferences may lead to significant metabolic impairment for only a small fraction of oxidoreductases. The widespread sequence conservation of NAD(H)/NADP(H) selective residues, however, seems to contradict this low sensitivity, suggesting that factors beyond flux stoichiometry may drive coenzyme redundancy. By developing a model of enzyme cost as a function of enzyme-coenzyme specificity we found that this conundrum can be resolved by viewing coenzyme redundancy as a graded improvement that enables efficient usage of protein resources when reactions are near thermodynamic equilibrium. These results suggest that widespread coenzyme redundancy in metabolic networks is the outcome of evolution for increased metabolic efficiency and parsimonious use of protein at the whole-cell level. Our findings may cast new light on the early stages and emergence cenzyme couplings, and could serve as design principles for the construction of modified or artificial metabolic circuits for synthetic biology and metabolic engineering applications.

## Results

### Thermodynamics constrain only a small number of NAD-coupled oxidoreductases

We first explored the robustness of metabolism to altering enzyme specificity of individual oxidoreductases to coenzymes as a way of investigating whether coenzyme coupling constrains metabolic networks (15). To investigate the consequences of altering the NAD(P)-specificity of individual oxidoreductases, we used flux balance analysis (FBA) (28) to simulate the growth rates of *in silico* mutants with altered coenzyme specificity in 117,180 different growth media. Specifically, we altered the iJO1366 (29) *E. coli* genome-scale metabolic model as follows: first, we changed stoichiometric coefficients to replace, one by one, the original coenzyme stoichiometric coefficients with the alternative coenzyme. In other words, if a reaction chosen for a mutation was originally coupled with NAD(H), it will now become coupled with NADP(H) instead. Since the *in vivo* NADH/NAD^+^ ratio is drastically different from the NADPH/NADP^+^ ratio (3, 8), some reactions that are reversible with the original coenzyme coupling may become irreversible upon switching coenzymes (Fig S2a-b), even when taking into consideration the possible range of concentrations of other metabolites in the cell (see Methods). Thus, upon switching coenzymes, we re-evaluated the thermodynamic feasibility of each reaction in either direction, and translated this information into constraints included into our altered metabolic model (Fig S2a-b, Methods). We compiled a list of 76 genes encoding NAD(P)-coupled oxidoreductases (Supplemental Table 1) and generated 76 single mutant metabolic models by switching the coenzyme preference of each enzyme, one at a time. For each model mutant, we then computed maximal growth rates in 117,180 media conditions, spanning fermentation, aerobic respiration and anaerobic respiration using nitrate, nitrite, dimethyl sulfoxide (DMSO), trimethylamine *N*-oxide (TMAO) or fumarate, on combinations of 180 carbon sources and 93 nitrogen sources (Methods, Supplemental Table 2).

We first counted the frequency of lethal mutations across all *in silico* media conditions (Fig. 2b, Methods), expecting coenzyme switching to cause major fitness defects for each gene under at least some conditions. To our surprise, the vast majority (97.3%, 8,103,217 of 8,323,596) of environment and coenzyme mutation pairs were non-lethal, suggesting that metabolism is overall robust to individual swaps of coenzyme specificity. However, of the 219,092 lethal mutant-environment pairs, all came from forcing one of six NAD(H)-coupled oxidoreductases to use NADP(H). These genes are: *leuB, pdxB, paaH, gatD, lgoD*, and *fucO* (Fig. 2b). By further constraining the range of allowed metabolite concentrations, we also found environments where sugar alcohol oxidoreductases encoded by *srlD* and *mltD* require coupling to NAD(H) rather than NADP(H) (Fig. S1). Only two oxidoreductases, encoded by the two genes *leuB* and *pdxB*, were predicted to require NAD(H) as a coenzyme rather than NADP(H) in all conditions tested (Fig. 2b). This is in agreement with extensive work by Dean and colleagues, who demonstrated that the gene product of *leuB*, isopropylmalate dehydrogenase, is strongly specific for NAD(H), and that introducing mutations to alter specificity towards NADP(H) always reduced fitness (13). Notably, all the observed lethal gene-environment pairs are alleviated when we do not take into consideration the thermodynamic differences between NAD(H) and NADP(H) (Fig. S2), demonstrating that the thermodynamic driving force is a dominant factor in constraining the coenzyme preference in our model.

**Figure 2:**
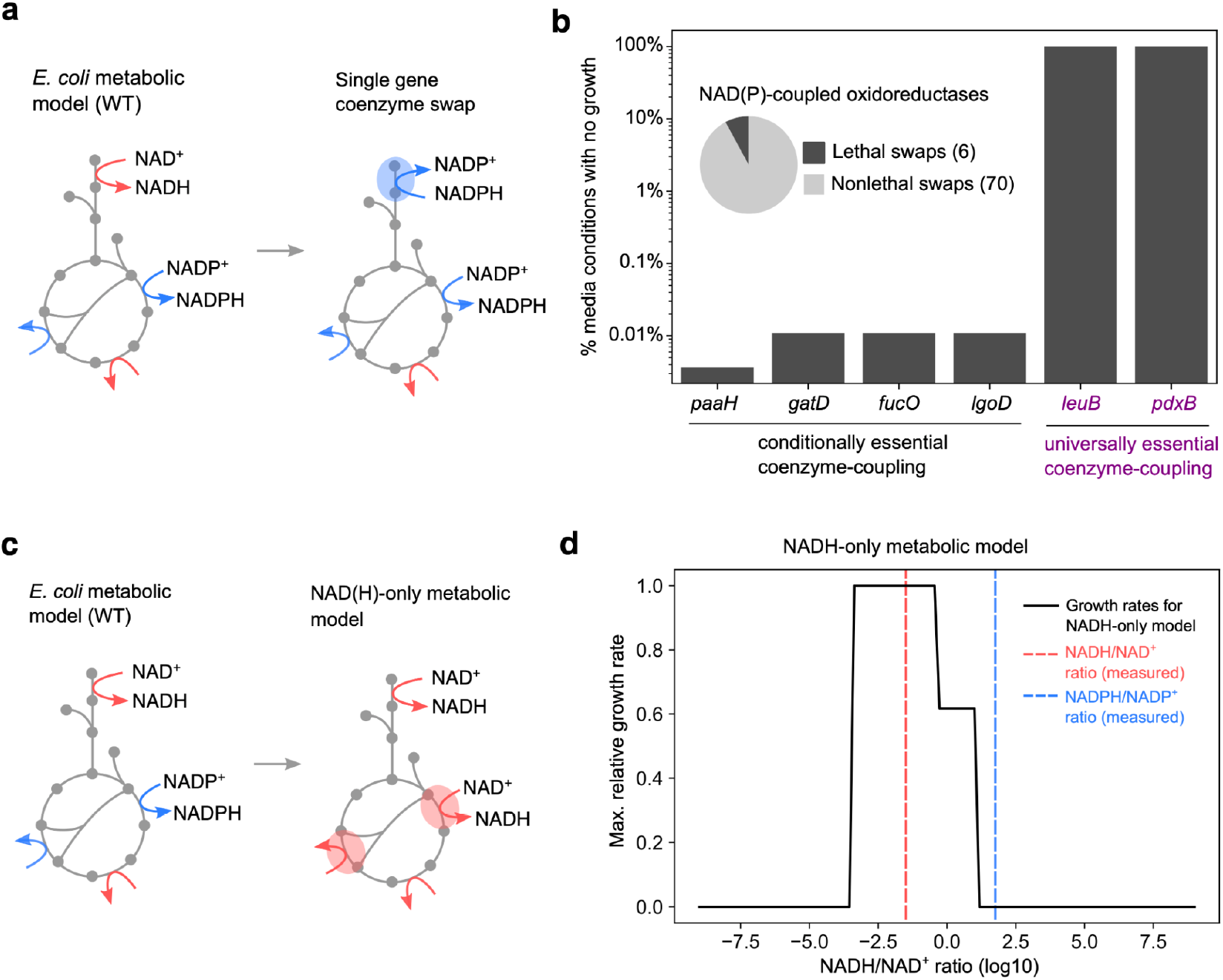
Coenzyme “rewiring” in genome-scale metabolic models. **a**. We determined the consequences of “rewiring” the *E. coli* metabolic network iJO1366 by changing the coenzyme specificity of 76 genes encoding NAD(P)-dependent oxidoreductases, and stimulated the growth across 117,180 growth media spanning different combinations of 180 carbon sources, 93 nitrogen sources and 6 electron acceptors (oxygen, nitrite, nitrate, TMAO, DMSO, fumarate) as well as fermentative growth using flux balance analysis (FBA). **b**. Only 6 of 76 genes were predicted to require a specific nicotinamide cofactor in one or more media conditions (*x*-axis, inset). The percentage of the media conditions that led to no growth is plotted as a bar graph (*y*-axis), showing that only two oxidoreductases are constrained to the endogenous coupling in all conditions (purple text): the NAD(H)-coupled isopropylmalate dehydrogenase *leuB* and erythronate 4-phosphate dehydrogenase *pdxB*. **c**. We next altered the *E. coli* metabolic model by replacing all NADP(H)-coupled reactions with NAD(H)-coupled reactions (see Methods), and simulated growth at various concentration ratios of NADH/NAD^+^. **d**. We computed the maximum growth rate of the NAD(H)-only metabolic model (*y*-axis) at various concentration ratios for NADH/NAD^+^ (*x*-axis, black line), and found that a wide range of ratios enabled growth at comparable levels to the unmodified model. The red and blue dashed line indicates the measured concentration ratios in *E. coli* for NADH/NAD^+^ (0.032) and NADPH/NADP^+^ (57.14), respectively. Note that a single coenzyme poised at the measured NADH/NAD^+^ ratio could enable cellular growth.

Our flux predictions suggest that enzymes that are sensitive to switching coenzymes are limited to NAD(H)-dependent, and not NADP(H)-dependent, oxidoreductases, suggesting that a metabolism using only NAD(H) might be feasible. To test this scenario, we constructed an NAD(H)-only model of *E. coli* metabolism by swapping all NADP(H)-coupled reactions with NAD(H), and removing NADP^+^ synthesis and degradation reactions (see Methods). We systematically computed maximum growth rates across a broad gradient of NADH/NAD^+^ ratios (Fig. 2c), and found that a wide range of NADH/NAD+ ratios enabled growths similar to that of the unmodified model (Fig. 2d). The feasibility of an NAD(H)-only *E. coli* metabolism is also corroborated by an alternative approach, i.e. thermodynamic flux balance analysis (TFBA), applied to a core *E. col*i model. (Fig. S3). Altogether, these results challenge the commonly accepted notion that two redox coenzymes are essential for running metabolism at the whole-cell level, and suggest that the reasons for coenzyme redundancy and NAD(H) or NADP(H) specificity might have to be sought elsewhere.

### Amino acid residues conferring selectivity to NAD(H) or NADP(H) are well conserved

Flux modeling predictions suggest that while the maximum organismal growth rate is insensitive to the coenzyme specificity for the majority of enzymes, a few genes are expected to be highly constrained to a single coenzyme specificity, limited to NAD(H)-coupled oxidoreductases. To determine if these model predictions are consistent with measures of enzyme specificity, we used a combination of protein structure and bioinformatic analysis to assess the coenzyme binding preferences of oxidoreductases both in *E. coli*, and across phylogenetically divergent species.

We first investigated *leuB* and *pdxB*, which both require NADH-coupling to permit biomass production and growth *in silico*. While *leuB* has been previously shown to use only NAD(H)(13), *pdxB*, which encodes the enzyme erythronate-4-phosphate dehydrogenase involved in *de novo* pyridoxal 5’-phosphate biosynthesis, has received less attention. Analysis of the structure of erythronate-4-phosphate dehydrogenase bound to NAD(H) in *Salmonella Enterica* indicates that an aspartic acid residue residue (Asp146) coordinates the 2’ and 3’ hydroxyl groups in the ribosyl portion of NAD(H), conferring selectively for NAD(H) over NADP(H) (Fig. S4) (30–32). We reasoned that if erythronate-4-phosphate dehydrogenase was constrained to use NAD(H) in all environments, then orthologues from diverse species living in heterogeneous environments should display similar specificity and, therefore, conservation of Asp146. We obtained all *pdxB* orthologs from the KEGG database (orthogroup K03473, *n*=1087), and performed multiple sequence alignment on the coenzyme-binding Rossman fold (Methods). We found that Asp146 was conserved in all *pdxB* orthologs, consistent with predictions that *pdxB* is universally required to use NAD(H) like *leuB* (Fig. 3a).

**Figure 3:**
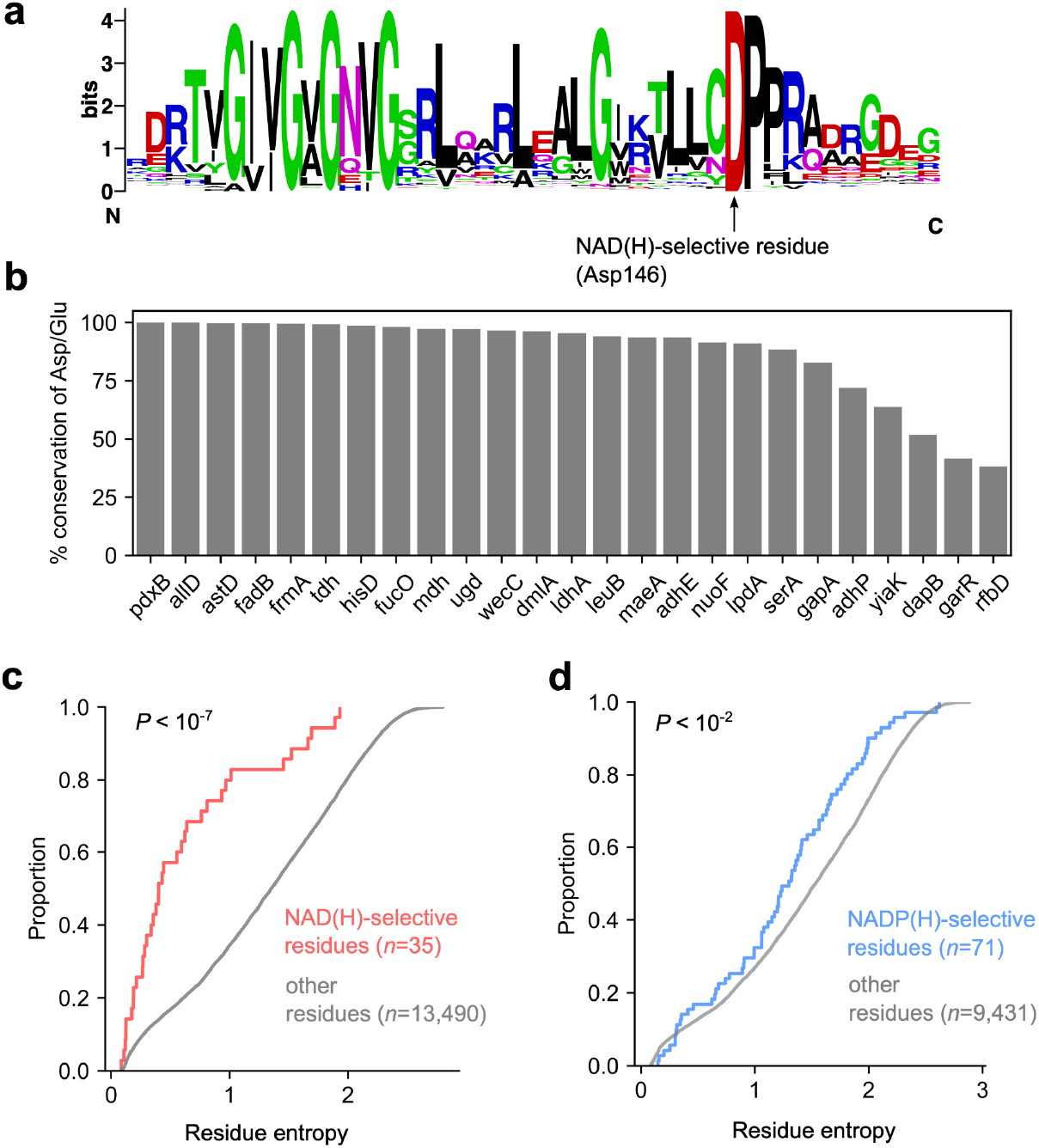
Flux balance models fail to fully explain the widespread conservation of coenzyme-selective residues in NAD(P)-coupled oxidoreductases. **a**. Orthologs of *pdxB* were retrieved from the KEGG database (K03473). The Rossman fold was identified using HMMR 3.1(47) to generate a sequence logo for the NAD(H) binding domain of these enzymes (Methods). The arrow denotes the 100% conservation of the aspartic acid residue (Asp146) responsible for discriminating between NAD(H) and NADP(H). **b**. Conservation of Asp or Glu residues coordinating hydroxyl groups in the ribosyl moiety of NAD(H) across 25 NAD(H)-coupled enzymes in the *E. coli* metabolic network. The *y*-axis is the percentage conservation of either Asp or Glu at the NAD(H)-selective site inferred from protein crystal structures (see Methods). **c**. We computed the entropy for each residue that conferred selectivity for either NAD(H) in 35 KEGG orthogroups encoding NAD-coupled oxidoreductases, and plotted the empirical cumulative distribution (eCDF) for residues that are responsible for NAD(H) selectivity (red) vs all other residues in the protein sequences (grey). **d**. The same as **c**., but for 25 NADP(H)-dependent oxidoreductases, where the entropy distribution for NADP(H)-selective residues are plotted in blue. NAD(H) and NADP(H) selective residues displayed significantly lower entropy than non-selective residues (Mann whitney U-test: *P* < 10^−6^ for NAD(H)-dependent oxidoreductases and *P* < 10^−2^ for NADP(H)-dependent oxidoreductases).

We next sought to determine whether the observed degree of conservation of Asp146 in *pdxB* orthologs is higher than other NAD(H)-dependent oxidoreductases that are not predicted to be constrained to use NAD(H). We determined whether there was structural evidence in the Protein Data Bank (PDB) for each NAD(P)-dependent oxidoreductase in the *E. coli* (or a homolog with greater than 30% homology) bound to NAD(P), and annotated residues that confer selectivity to NAD(H) or NADP(H). We found 25 oxidoreductases with NAD(H)-bound structures in the PDB with clear evidence of hydrogen bonding between an aspartic acid or glutamic acid residue and the 2’ and 3’ hydroxyl groups in the ribosyl portion of NAD(H) (Supplemental Table 3). We performed multiple sequence alignment for orthogroups for each oxidoreductase, and quantified the conservation of the aspartic acid or glutamic acid residue across orthologs (Methods, Fig. 3b). While the universally constrained oxidoreductase encoding genes *leuB* and *pdxB* show extremely high conservation of the coordinating aspartic acid residue (94 and 100%, respectively), several other oxidoreductases are not predicted to be constrained to using NAD(H) show comparably high conservation of the coordinating aspartic acid or glutamic acid residue (Fig. 3b). More broadly, analysis of residues that confer selectivity for NAD(H) or NADP(H) showed that these residues were significantly less variable than the remaining portion of the protein sequences (Fig 3c-d), and the degree of conservation is not associated with flux balance predictions.

Altogether, the results from genome-scale simulations suggest that metabolism, in principle, can operate with a single nicotinamide coenzyme system. On the other hand, sequence conservation patterns indicate clear conserved preferences for specific coenzyme pairings for a number of enzymes. One possible explanation for this apparent discrepancy is that coenzyme specificity may be selectively advantageous without being essential to carry out metabolic functions, and that a two-coenzyme system may have a quantifiable fitness advantage over a metabolic network with a single-coenzyme.

### Enzyme cost minimization drives separation of coenzyme preference globally

Thus far, the fitness consequences of modifying oxidoreductase specificity for different coenzymes has been assessed by computing maximum growth rates using stoichiometric models of metabolism with altered coenzyme couplings for individual enzymes. These models suggest that thermodynamics may drive specificity for a few individual enzymes, giving rise to “local”‘ constraints for enzyme specificity that are also recapitulated in protein sequence-level variation. However, these models also suggest that a metabolism with a single coenzyme (poised close to the in *vivo* NADH/NAD^+^ ratio) can satisfy these constraints, which is in stark contrast with the observed widespread specificity for NAD(H) or NADP(H) by oxidoreductases (Fig. 3).

Beyond inducing local constraints in single reactions, thermodynamics may also induce “global” fitness consequences through the control of the total protein demand to carry biosynthetic flux. This is due to the fact that strong thermodynamic driving forces in enzyme-catalyzed reactions reduce the minimum protein abundance necessary to maintain a fixed reaction flux (33–36). This effect comes directly from the following flux-force relation (37):

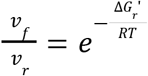

where Δ*G*_*r*_′is the free energy at physiological metabolite concentrations, and *v*_*f*_ and *v*_*r*_ is the forward and reverse reaction flux, respectively. Note that as the free energy decreases, the proportion of flux that is in the forward direction increases, and thus decreases the enzyme abundance required for catalysis (33, 34). We hypothesized that coenzyme redundancy might allow the cell to lower the total protein required to maintain intracellular reaction fluxes. As in previous work, we used the flux-force relation to model the interrelationship between thermodynamics and enzyme cost(33–36). Specifically, we constructed a model of enzyme cost as a function of coenzyme specificity, and aimed to study how the number of coenzymes affects the amount of protein enzymes needed to run metabolism (Fig. 4a, Methods).

**Figure 4:**
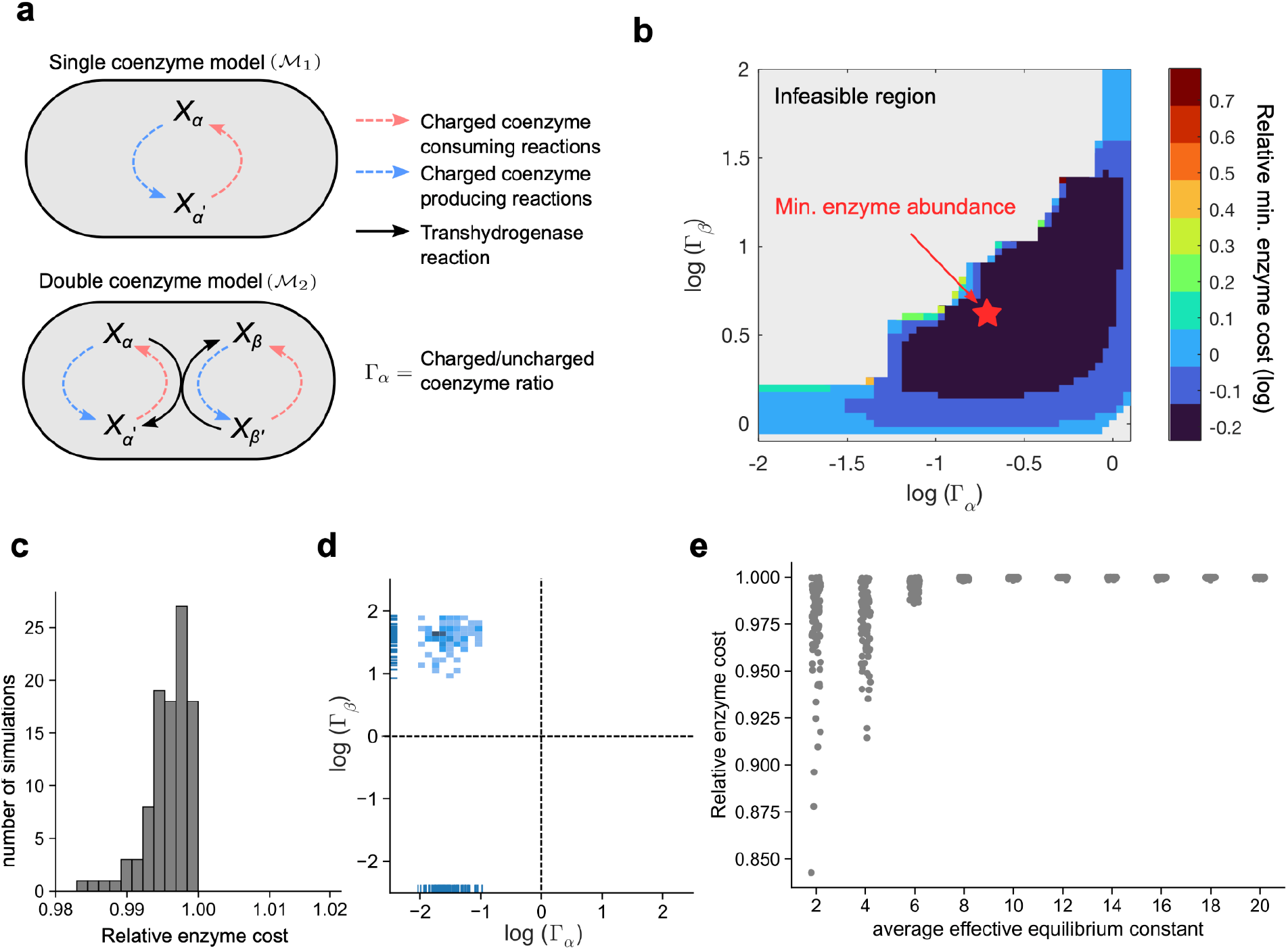
A minimal model of coenzyme-utilizing enzyme cost predicts the emergence of coenzyme redundancy. **a**. Model of a reaction network where multiple reactions (dashed arrows) produce and consume a charged coenzyme, where reaction flux is feasible with a single coenzyme (top), compared to a model where reaction flux is partitioned between two coenzyme pools connected by a transhydrogenase reaction (bottom). **b**. A heatmap of the minimum enzyme cost (scaled by the enzyme cost for a single-coenzyme model) as a function of various coenzyme ratios (*x* and *y* axis) for randomly sampled reaction fluxes, thermodynamic and kinetic parameters (see Methods). The color scale denotes the minimum relative enzyme cost at each coenzyme ratio (log-scale), and the grey region denotes the region of space that is thermodynamically infeasible. Note that the dark blue regions denote the space of coenzyme ratios that enables enzyme cost to be less than the single coenzyme, and the red star denotes the point where the minimum enzyme abundance is achieved. **c**. We simulated 100 random realizations, and plotted the distribution of the minimal enzyme cost relative to the cost using a single coenzyme system (*x*-axis), and found that all simulations were lower than unity. **d**. The coenzyme ratio for coenzyme α(*x*-axis) vs coenzyme β(*y*-axis) that minimizes total protein abundance were plotted for each realization as a 2-D histogram. **e**. We varied the mean effective equilibrium constant (μ_*K*_, *x*-axis), and plotted the fractional reduction in minimal protein required (*y*-axis) relative to a single coenzyme model (see Supplementary text for definitions and derivations).

Given the uncertainty about many model parameters, we generated predictions based on extensive random sampling of key parameters, including the reaction network topology, the effective equilibrium constant for each reaction *r* (*K*^†^_*r*_), the maximal turnover rates (κ_*r*_), and the flux through each reaction (*v*_*r*_). In this formulation, the effective equilibrium constant aggregates the effects of the free energy change at standard molar conditions, and the concentration differences between products and substrates that are not coenzymes, capturing the coenzyme-independent forces driving reaction *r* (see Supplementary Text). Additionally, our model also relies on quantities related to the thermodynamic drive of each coenzyme as well as the flux partitioning between different coenzyme pools. There features are captured by the ratio of charged to uncharged species for various coenzyme pairs 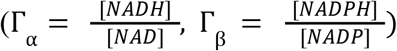 and the fraction of flux for reaction *r* through the NAD(H) coenzyme pool (*v*_*r*α_) and NADP(H) pool (*v*_*r*β_). We next used nonlinear optimization techniques to find the coenzyme concentration ratios (Γ_α_, Γ_β_) and the reaction fluxes through each coenzyme pool(*v*_*r*α_, *v*_*r*β_) that minimized the ; total protein abundance in the network (Fig. 4b, see Supplementary text). We found that partitioning flux through two coenzyme pools decreased the minimum protein abundance required relative to the single coenzyme scenario (Fig. 4b). Notably, at the optimum, each reaction flux was driven through primarily one coenzyme, indicating that enzyme specificity for a single coenzyme could emerge from the cellular-level objective of optimizing proteome allocation (Fig. S5a). The emergence of enzyme specificity for one of the two coenzymes was driven by the combination of both kinetic and thermodynamic factors (Fig. S5b-c), where reactions that required less protein to maintain the given flux demand were used to re-balance coenzyme pools, operating against the coenzyme concentration gradient (Fig. S5b-c).

To see if these results were generic for different realizations of parameters, we computed optimal flux distributions and coenzyme ratios for 100 random instances. The two-coenzyme model always achieved lower total protein cost than the one coenzyme model (Fig. 4c). For each simulation, we plotted the coenzyme ratios found to minimize the total protein abundance, and we found that all simulations resulted in optimal ratio sets with opposing thermodynamic drive, where one coenzyme pool was primarily in the charged state, and the other was primarily in the uncharged state (Fig. 4d). This is strikingly consistent with the observation that *in vivo* measured ratios of oxidized to reduced NAD(H) and NADP(H) coenzymes are greater than and less than one, respectively (3, 8). Interestingly, our model also predicts that the largest reduction in enzyme cost comes when the mean effective equilibrium constants are closer to unity (Fig. 4e), suggesting that two-coenzyme systems may be particularly beneficial if the coupled reactions are held closer to equilibrium.

### NAD(P)-dependent oxidoreductase coenzyme-specificity is associated with reaction thermodynamics

From the models presented above, we found that enzyme preference for specific coenzymes is strongly associated with thermodynamic drive, where enzymes catalyzing reactions near equilibrium are predicted to be more specific for one coenzyme over another (Fig. 5a). To test this prediction, we computed free energies for all NAD(P)-coupled oxidoreductases in the KEGG database using eQuilibrator(38, 39), and computed binding preferences of protein sequences for NAD(H) and NADP(H) using an artificial neural network (ANN)(40), which estimates the likelihood of FAD(H2), NAD(H) and NADP(H) binding in protein sequences containing Rossman folds. We noticed that the distribution of free energies for NAD(P)-coupled oxidoreductases were bimodal, where oxidoreductases using oxygen as a co-substrate were driven far from equilibrium (Δ*G*_*r*_^′^° =421.8 +/-155 kJ/mol) compared to oxygen-independent oxidoreductases (Δ*G*_*r*_^′^° =12.4 +/-48.5 kJ/mol, Fig. 5b), providing a simple way to categorize oxidoreductases by thermodynamic potential. We computed binding preferences for NAD(H) and NADP(H) (Fig. 5c), and computed the proportion of Rossman folds that were single coenzyme binders (e.g., NAD(H) or NADP(H)) or multi-coenzyme binders (e.g., NAD(H) and NADP(H)). Consistent with predictions from the theory, we found that 61% (2139/3497) of folds from oxygen-coupled NAD(P)-dependent oxidoreductases are expected to be multi-coenzyme binders, compared to just 23% (28992/126718) from non-oxygen coupled NAD(P)dependent oxidoreductases (Fisher’s exact test: *P* < 10^−5^).

**Figure 5:**
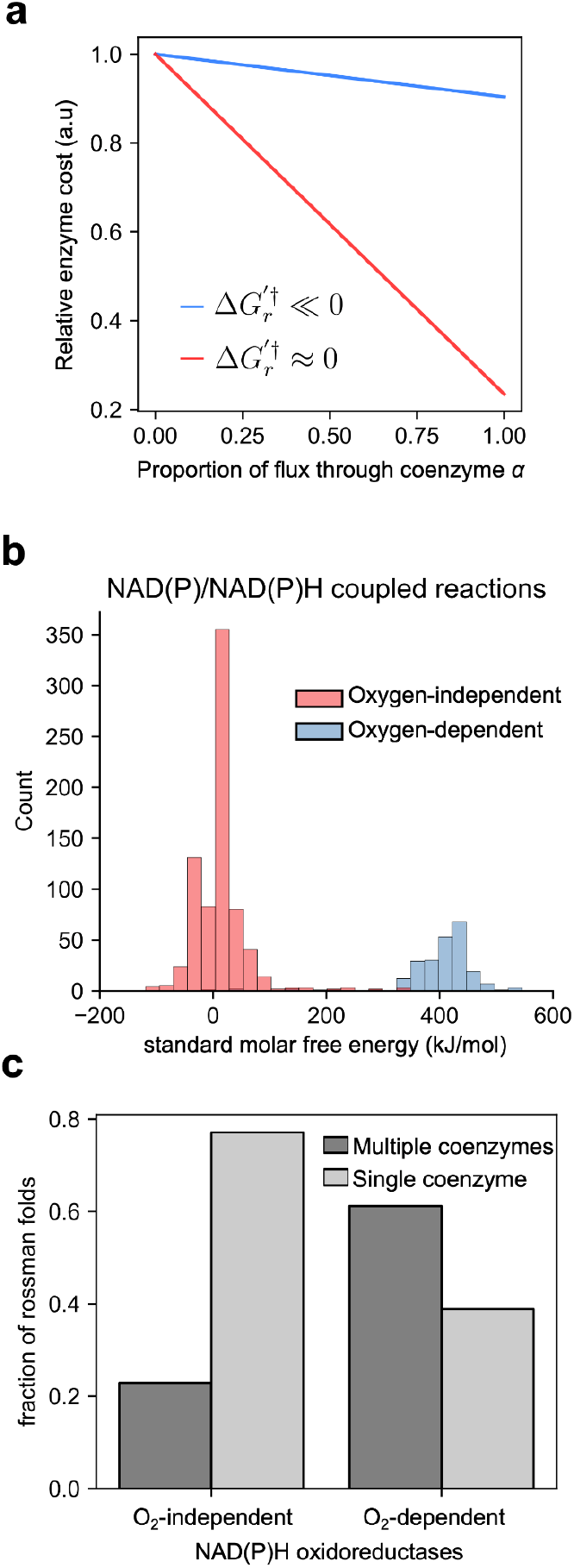
Oxygen-coupled NAD(P)-oxidoreductases are predicted to be more coenzyme promiscuous. **a**. The relationship between the degree of flux partitioning between two coenzyme pools (*x*-axis) and enzyme cost (*y*-axis, re-scaled by the maximum enzyme cost per model) at both low (red) and high (blue) coenzyme-independent driving forces (Δ*G*_*r*_′^†^). Note that at high driving forces, the sensitivity of enzyme cost to flux partitioning is significantly less than at low driving forces. **b**. We computed standard molar free energies for all NAD(P)-coupled reactions in the KEGG database using eQuilibrator (39), and plotted the distribution of free energies for (a) NAD(P)-coupled reactions in the NAD(P)H producing direction (*n*=990 reactions). Distributions of free energies from oxidoreductase reactions that use molecular oxygen as a co-substrate are plotted in blue, while distributions of free energies for other NAD(P)-coupled reactions are plotted in red. **c**. The proportion of Rossman folds that were predicted using an artificial neural network model (40) to bind a single coenzyme (light grey bars, e.g., NAD(H) or NADP(H)) or multiple coenzymes (dark grey bars, e.g., NAD(H) and NADP(H)) are plotted for oxygen-dependent or independent NAD(H)/NADP(H) oxidoreductases (*x*-axis). We found 61% of the folds from oxygen-dependent NAD(H)/NADP(H) oxidoreductases (2139/3497) were multi-coenzyme binders, compared to just 23% amongst oxygen-dependent NAD(H)/NADP(H) oxidoreductases (28992/126718) (Fisher’s exact test: *P* < 10^−5^).

### Dephospho-CoA might be a redundant coenzyme for acyl transfer reactions

The simple phenomenological model presented above (Fig. 4) demonstrates that coenzyme redundancy (i.e. multiple coenzymes for the same group transfer reaction) may be a generic mechanism to reduce intracellular protein cost, potentially explaining the maintenance of both NAD(H) and NADP(H) as intracellular redox coenzymes. However, this simple model is agnostic to the chemical details of the group or electron transfer (apart from thermodynamic potentials), suggesting that protein cost can be minimized by simply adding additional coenzymes to the intracellular repertoire. This begs the question of why several coenzymes appear to exist as only one variant in the cell.

One interesting example is Coenzyme A (CoA), a molecule involved in the redistribution of intracellular acyl groups. We computed free energies for various group transfer reactions (Fig. 6a), and found that the distribution of free energies for acyl transfer reactions were similar to NAD(P)-coupled reactions, suggesting multiple coenzyme systems may be also advantageous for CoA-coupled acetyl-transfer reactions. Interestingly, the last step of CoA biosynthesis resembles the last step of NADP biosynthesis, where a kinase acts on the ribosyl moiety of the coenzyme structure (Fig. 6b). For the case of NADP^+^ biosynthesis, NAD kinase phosphorylates NAD^+^ at the 3’ hydroxyl of the ribosyl moiety to create NADP^+^. For the case of CoA, dephospho-CoA kinase phosphorylates the 2’ hydroxyl of the ribosyl moiety of dephospho-CoA to produce CoA. The strikingly similar reaction motif may indicate that the dephospho-CoA is an active and alternative coenzyme capable of performing similar functions as CoA, analogous to NAD^+^ and NADP^+^. Interestingly, mass spectrometry experiments have shown that dephospho-CoA can be acylated in rat liver (41). We thus propose that dephospho-CoA may be a redundant coenzyme for acyl-transfer reactions rather than simply serve as an intermediate during CoA biosynthesis.

**Figure 6:**
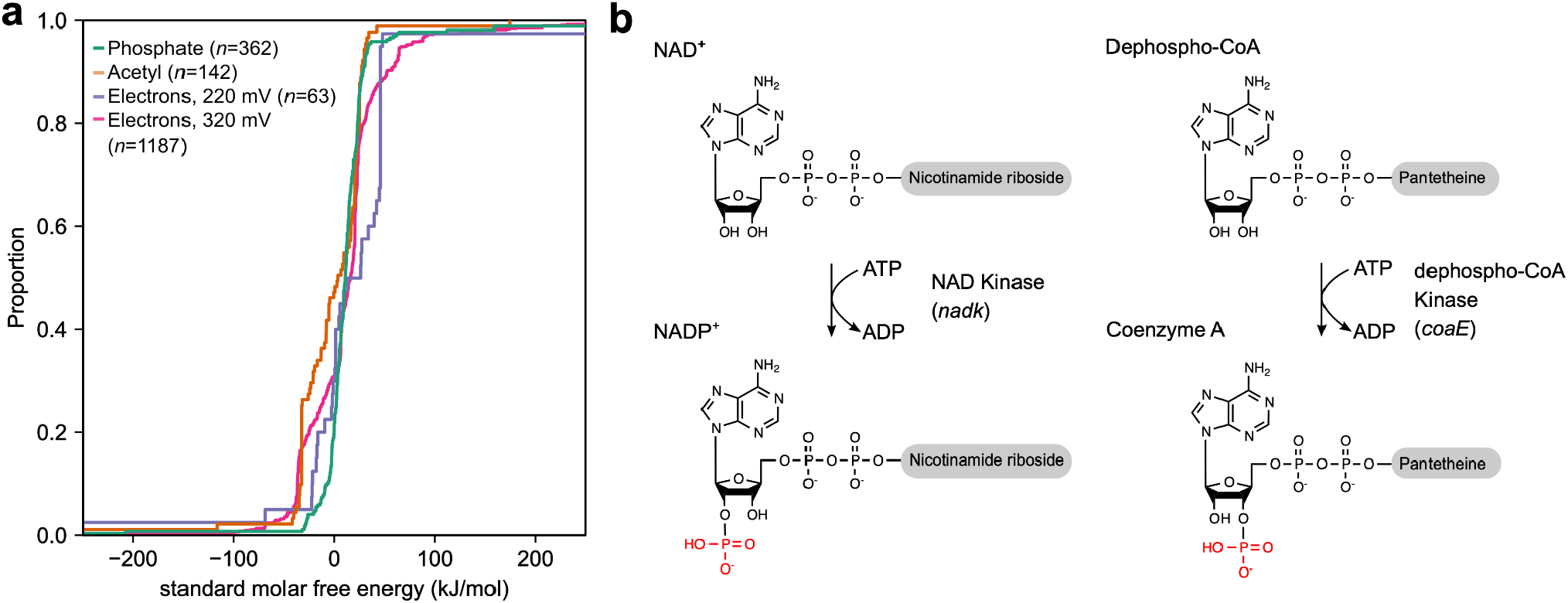
Comparison between NAD(P) and Coenzyme A biosynthesis reveals potential redundancy for acyl-CoA’s. **a**. Free energies for group transfer reactions of phosphate groups, acetyl groups and electron transfer reactions in KEGG were computed using eQuilibrator (38, 39), and the empirical cumulative distributions across reactions were plotted. Free energies for all group and electron transfer reactions are similarly distributed near equilibrium, suggesting that Coenzyme A-coupled reactions may also depend on redundant coenzyme systems similar to NAD(H)/NADP(H) . Free energies for redox and group transfer reactions were computed for the following coenzyme pairs. Phosphate-coupled reactions: ADP/ATP, CDP/CTP, GDP/GTP; Acetyl-coupled reactions: Coenzyme A/Acetyl-CoA; Electrons (220 mV): FAD/FADH2, FMN/FMNH2, Glutathione disulfide/Glutathione; Electrons (320 mV): NAD/NADH, NADP/NADPH. **b**. The last biosynthetic step of NADP^+^ synthesis is the phosphorylation of the 3’ hydroxyl group on the ribose moiety of NAD^+^, compared to the biosynthesis of Coenzyme A, where the 2’ hydroxyl of the ribose moiety is phosphorylated by dephospho-CoA kinase (EC 2.7.1.24)

## Discussion

The evolution of coenzyme coupling in redox biochemistry constitutes a fascinating puzzle central to the emergence and complexification of life itself (24, 42). In order to understand the constraints and the selection pressure that led to the enzyme-coenzyme couplings observed today, one would ideally shuffle all of these couplings (e.g., by reassigning coenzymes to different enzymes), and determine fitness changes under many different environments. While such an endeavor would be very challenging experimentally, it can be pursued efficiently using genome scale stoichiometric models of metabolism. In particular, we used flux balance modeling to survey dozens of *in silico* mutants of oxidoreductase enzymes in thousands of environmental conditions, explicitly modeling the consequences of changing an enzyme’s coenzyme preference on growth rate. Our analysis suggests that few enzymes are universally constrained to use NAD(H) or NADP(H), and that most dependencies are only observable in specific environments. Future experimental work could test these predictions by altering coenzyme specificity for some of these oxidoreductases and measuring growth rate in various media conditions (13).

Our flux modeling results suggest that thermodynamic constraints play a role in shaping the preferences of just a small number of oxidoreductases for specific coenzymes. Structural and bioinformatics analyses of *pdxB* suggests that these thermodynamic constraints can create strong selective pressures shaping protein sequence evolution, evidenced by our observation that all *pdxB* orthologs contain a conserved residue that specifically coordinates NAD(H), and potentially sterically prevents the binding of NADP(H) (Fig. 3a). However, our results also suggest that residues conferring selectivity for NAD(H) or NADP(H) in oxidoreductases are well conserved (Fig. 3b), indicating that constraint-based modeling may not adequately capture the principal factors driving enzyme specificity for individual coenzymes.

The model presented in Fig. 4 suggests that the emergence of widespread enzyme specificity towards one of many coenzymes could be induced by a selective pressure to minimize the total abundance of enzymes in the cell. Importantly, our model does not rely on chemical details specific to nicotinamide coenzymes or electron transfer, and could potentially explain the ubiquity of multi-coenzyme systems (e.g. ferredoxins, thioredoxins, glutaredoxins, flavins, quinones, hemes, pterins, nucleotide phosphates) in cellular metabolism. Notably, this model suggests that enzyme specificity towards one of the two coenzymes may be a generic strategy to reduce the total protein cost of oxidoreductase enzymes, and that coenzyme choice is shaped by both kinetic and thermodynamic factors. Additionally, feasibility of an NAD(H)-only metabolism supports the hypothesis that ancient metabolic networks may have originally relied on NAD(H) only, and gradually evolved NADP(H) dependencies. While it has been proposed that an NADP(H)-dependent isocitrate dehydrogenase (IDH) evolved from an ancestral NAD(H)-dependent enzyme in prokaryotes (14), future efforts to reconstruct ancestral sequences from various NAD(P)-dependent oxidoreductases can explore whether this is a general feature of NADP-coupled enzymes (43).

The flux analysis presented in this study only considers stoichiometric and thermodynamic factors shaping cell fitness. However, other constraints or metabolic demands exist that may constrain oxidoreductase coenzyme preference. For example, our flux models may not accurately capture metabolic demands induced by oxidative stress, as NADPH is the primary electron donor for enzymatic detoxification of reactive oxygen species and other damaging oxidants (2). Specificity may be shaped through mechanisms not directly linked to stoichiometric balancing or thermodynamics, including epistatic mechanisms, where the sequence variation in the coenzyme binding site indirectly influences co-substrate specificity, kinetics(44) or thermostability(45). Nonetheless, our results suggest that thermodynamics and enzyme cost minimization are important factors that can constrain the oxidoreductase specificity for NAD(H) or NADP(H).

In addition to offering an explanation for the occurrence of multi-coenzyme systems, our model can be used for the targeted search for unexplored redundant coenzymes. Our results point to the intriguing possibility that dephospho-CoA might serve as an acyl-transfer coenzyme rather than simply an intermediate during CoA, and could in principle explain the occurrence of acyl-dephospho-CoAs(41). More broadly, the principle that enzyme cost minimization universally promotes coenzyme redundancy may help guide the interpretation of molecular diversity in the biosphere (46), as well as serve as a design principle for metabolic engineering and synthetic biology applications.

## Acknowledgements

We thank Arren Bar-Even, Adrian Jinich, Igor Libourel and Pankaj Mehta for helpful discussions. We acknowledge the support provided by the Directorates for Biological Sciences (BIO) and Geosciences (GEO) at the NSF and NASA under Agreements No. 80NSSC17K0295, 80NSSC17K0296 and 1724150 issued through the Astrobiology Program of the Science Mission Directorate. J.E.G. is supported by the Gordon and Betty Moore Foundation as Physics of Living Systems Fellows through grant number GBMF4513.

## Contributions

J.E.G.. and D.S. designed the research. J.E.G. wrote code and ran simulations. J.E.G. and A.I.F. performed analysis. A.B.G. contributed to protein cost modeling. J.E.G., A.I.F., and D.S. wrote the manuscript. All authors read and approved the final manuscript.

## Competing Financial Interests

The authors declare no competing financial interests

## Materials and Methods

### Software availability

Code and data are available on the following github repository: https://github.com/jgoldford/coenzymes

### Metabolic modeling of oxidoreductase specificity

We computationally accessed the fitness consequences of rewiring the NAD(P)H specificity of individual metabolic genes in *E. coli* by altering stoichiometric and thermodynamic constraints in a genome-scale metabolic model (GEMM), and by simulating the growth rate under various media conditions. We used the GEMM iJO1366(29) downloaded from the BIGG database (48), with Gibbs Free Energies derived using the component contribution method (49), using the eQuilibraitor API at pH 7 and ionic strength 0.1 M(38, 39). For each oxidoreductase gene that mapped to a reaction in iJO1366, we altered the coenzyme preference by first changing the stoichiometric coefficient in that reaction to the alternative coenzyme. For genes encoding reactions that utilized both coenzymes, we altered specificity by forcing all reactions to use only NAD(H) or NADP(H). To model the thermodynamic effect of changing coenzymes specificity, we used the computed reaction free energies at standard molar conditions, and bounds on intracellular metabolite concentrations to estimate the maximum and minimum driving force obtainable for each reaction using the following equation:

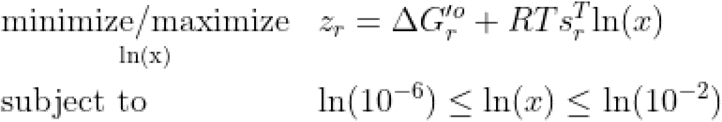

where *s*_*r*_ is the vector of stoichiometric coefficients for reaction *r*, Δ*G*_*r*_ ^′^° is the change in free energy at standard molar conditions for reaction *r, R* is the gas constant, *T* is temperature in Kelvin and *ln*(*x*)denotes the vector of natural log-transformed metabolite concentrations.. Note that we fixed the concentration of coenzymes based on previously measurements, where [NAD] = 2. 6 × 10^−3^, [NADH] = 8. 3 × 10^−5^, [NADP] = 2. 1 × 10 ^−6^, and [NADPH] = 1. 2 × 10^−4^ . We then allowed all other metabolite concentrations to vary when predicting the maximum and minimum driving forces for each reaction. After each *in silico* mutation that changed coenzyme preferences, we re-computed the minimum and maximum driving force, and used these values to set the upper and lower bounds of reactions, such that:

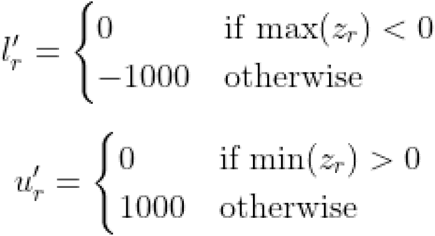

Where *l*′_*r*_ and *u*′_*r*_ are the updated lower and upper bounds for reaction *r*, respectively. We simulated the consequences of changing oxidoreductase enzyme specificity for NAD(P)H in *E. coli* by then simulating the maximum growth rate using flux balance analysis(28) under 117,180 conditions, consisting of various carbon sources, nitrogen sources and electron acceptors (Oxygen, Nitrite, Nitrate, TMAO, DMSO, fumarate) as well as fermentative growth, using the following linear program:

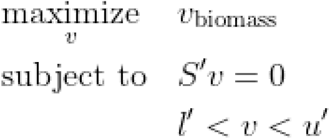

Where *S*′, *l*′, *u*′ is the altered stoichiometric matrix, flux lower bound, and flux upper bound after updating stoichiometric and thermodynamic constraints. Note that for the NAD(H)-only metabolism presented in Fig. 2c-d, we performed the same procedure described above but for all NADPH-coupled reactions, and under a range of different coenzyme ratios (Fig. 3c-d). For these simulations, we removed NAD kinase (NADK), NAD phosphatase (NADPPPs), NAD diphosphatase (NADDP), and NAD transhydrogenase (NADTRHD) from the model, and performed FBA in aerobic conditions with glucose as the sole carbon source.

For results presented in Fig. 2b, we restricted the analysis to the media sets (109,521 conditions) where growth of the unperturbed network was feasible. We modeled mutations for 76 genes encoding oxidoreductases in *E. coli*. This set of genes were determined using the following criteria: (a) the gene encoded an enzyme that could be classified using the following enzyme commission code - E.C. 1.X.1.Z, where X was any number except 6, and Z= any number, and (b) there were no other genes coding for enzymes that catalyze the same reaction. All simulations were performed using CobraPy and Gurobi (version 9.0.0) optimizer (50).

Thermodynamic flux balance analysis was performed using pyTMFA (51), with the reduced core *E. coli* metabolic model irJO1366 (52). We modified irJO1366 by substituting all NADP(H)-coupled reactions with NAD(H)-coupled reactions, and removed the NAD(P)H transhydrogenase reaction, and performed TMFA as previously described (51).

### Oxidoreductase specificity from compiled structural data and sequences

We compiled a list of NAD(P)-dependent oxidoreductases in the *E. coli* genome, and manually searched for structural features that confer selectivity of NAD(H) vs NADP(H). To this end, we identified orthogroups (KO) using the KEGG REST API, and developed a custom python script to identify NAD(P)-bound protein data bank (PDB) codes from KEGG genes within each orthogroup. For each orthogroup that contained at-least one NAD(P)-bound structure for the *E. coli* ortholog, or an ortholog with at-least 30% homology, we manually identified residues that clearly showed hydrogen bonding with the 2’ and 3’ hydroxyl group of the ribosyl moiety in NAD(H), or electrostatic interactions with the 3’ phosphate in NADP(H).

We first analyzed the conservation of the residues conferring specificity in orthologs of *pdxB* (K03473), the gene that codes for erythronate-4-phosphate dehydrogenase. We downloaded all orthogroups in the KEGG database (orthogroup K03473, *n*=1087), and performed multiple sequence alignment on the coenzyme-binding Rossman fold. We first identified a 42 amino acid subsequence in the *E. coli* genome using HMMR 3.1 as the Rossman fold in the *pdxB* gene (b2320) from *E. coli* MG1655, and used the biopython “pairwise2” module to perform local alignment between the reference Rossman fold and the ortholog sequence. Parameters were chosen such that gap opening and extension were penalized much more than non-matching characters (matched character: 1; unmatched character: 0; gap opening penalty: -10; gap extension penalty: -1). For each ortholog, we identified the subsequence with the largest score and labeled this subsequence as the Rossman fold for each ortholog. The sequences were then aligned using MUSCLE v3.8.31 using default parameters.

To compare other genes to *pdxB*, we identified 25 genes encoding enzymes in the iJO1366 model with structural evidence supporting the coordination of a single aspartic acid or glutamic acid residue and the 2’ and 3’ hydroxyl group of the ribosyl moiety in NAD(H) (Supplemental Table 3). For orthogroups with >1000 orthologs, we downsampled the sequences to 1000 before identifying running multiple sequence alignment.

We annotated residues that confer coenzyme selectivity for 57 NAD(P)-coupled oxidoreductases in *E. coli* based on structures available in the PDB (Supplementary Table 3). We performed multiple sequence alignment using MUSCLE, and computed the shannon entropy, *H*_*i*_, at each residue position *j*, and classified a residue as conferring selectivity to NAD(H) vs. NADP(H), such that:

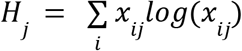

and *x*_*ij*_is the proportion of sequences with amino acids residue *i* in position *j*. Prior to computing distributions of entropy across residues that confer selectivity to either NADH or NADPH, we removed all residues with >50% gaps, as these represented poorly aligned regions of the protein sequence.

In Fig. 5c and Fig. S6, we computed coenzyme binding likelihoods using an artificial neural network model called Cofactory (40). Briefly, Cofactory first identifies the Rossman folds in a protein sequence using HMMR 3.1, and estimates the likelihood each fold binds FAD(H2), NADH and NADPH. We identified 779 orthogroups that either use oxygen as a co-substrate, and downloaded all sequences within each orthogroup from the KEGG REST API (53). Like in the previous case, we downsampled the sequences to 1000 if the orthogroup contained more than 1000 sequences before running Cofactory. After obtaining predictions of binding likelihoods to NAD(H), NADP(H) and FAD(H2), we filtered out Rossman folds that either (a) unambiguously bound FAD(H2) or (b) were not predicted to bind any of the coenzymes. We then categorized the remaining Rossman folds as “multi-coenzyme binders’’ if the binding likelihood exceeded 0.5 for more than one coenzyme, and “single coenzyme binders” if the binding likelihood exceeded 0.5 for either NAD(H) or NADP(H), but not both.

### Phenomenological modeling of proteome cost for multi-coenzyme systems

A detailed derivation of the phenomenological model used to analyze protein cost for multi-coenzyme systems in Fig. 4 is provided as a Supplementary Text. Here we present the final model used in the main text. The following non-linear optimization problem was constructed to find the minimal protein cost by varying coenzyme concentration ratios and flux partitioning, such that:

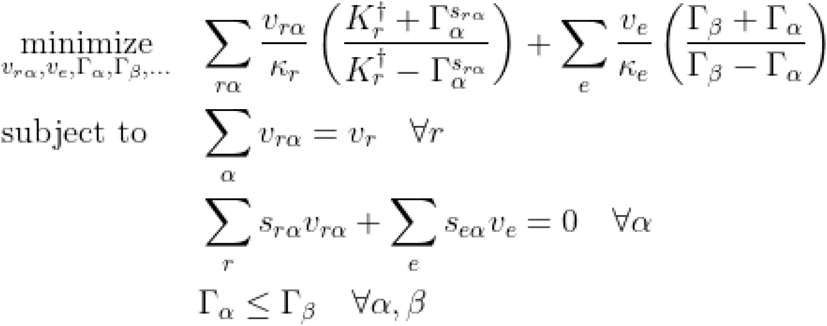

where sampled parameters are the following: *K*^†^_*r*_ is the effective equilibrium constant for reaction *r*, κ_*r*_ is the maximum turnover rate for reaction *r, v*_*r*_ is the net flux for reaction *r*, and *s*_*r*α_ is the stoichiometric coefficient for the charged coenzyme (negative if the charged-coenzyme is being consumed and is positive if being produced). Variables in our optimization approach are the following: Γ_α_the ratio of charged to uncharged species for coenzyme pair α, *v*_*r*α_ is the fraction of flux from reaction *r* through the coenzyme pool α. In this model, we also include an additional set of reactions *e*, which are generalized exchange reactions that shuttle groups between coenzymes, much like NAD(P)H transhydrogenase enzymes. The objective function is the total enzyme cost of the coenzyme-coupled sub-network plus the cost of all generalized exchange reactions (see Supplemental Text for derivation). All enzyme cost simulations were performed in MATLAB R2021a, using the COBRA toolbox (50) and GLPK solver for linear programs.

## Notes

### Competing Interest Statement

The authors have declared no competing interest.

