## Supplementary Text for "Protein cost minimization promotes the emergence of coenzyme redundancy"

### Supporting Information Text

In this supplemental note, we provide a detailed explanation for the model presented in Fig. 4a in the main text, where we model the enzyme abundance requirement to catalyze flux distributions using a single coenzyme or two coenzymes. Prior studies have shown that the amount of enzyme required to catalyze a net flux increases exponentially when decreasing thermodynamic drive (1–3). Thermodynamic flux balance models (4) do not capture this quantitative relationship between driving forces and enzyme abundance, but rather simply capture the qualitative requirement that if a reaction carries net positive flux, $v_r > 0$, then the change in the Gibbs Free Energy must be negative, such that $\Delta G_r < 0$. The goal of developing the forthcoming model is to explore the potential quantitative relationship between coenzyme preference/diversity and enzyme abundance.

**Enzyme cost model with multiple coenzymes.** Suppose that reaction $r$ can be catalyzed by an enzyme that utilizes one of many coenzyme pairs, (denoted by the greek index $\alpha$), such that an enzyme (with abundance $E_{r\alpha}$) carries out the following reaction scheme:

$$S_r + \alpha \longrightarrow P_r + \alpha'$$

where $S$ and $P$ represent the primary substrates and products of reaction $r$, respectively, and $\alpha$ represents the coenzyme in the uncharged state (e.g. NAD(P)$^+$) and $\alpha'$ represents the coenzyme in the charged state (e.g. NAD(P)H). The driving force for this reaction can be represented using the following expression:

$$\Delta G'_r = \Delta G'^{\circ}_r + RT\ln\frac{x_p x_{\alpha'}}{x_s x_\alpha}$$

where $x_p, x_s, x_{\alpha'}, x_\alpha$ represent the activities of substrate, product, charged coenzyme, and uncharged coenzyme, respectively. Note that we can separate the driving force contributions from the substrates and products and coenzymes, and aggregate the driving force contributions from the free energy change at standard molar condition with the substrate and product driving force, such that $\Delta G^{\dagger}{}'_r = \Delta G^{\circ}{}'_r + RT\ln\frac{x_p}{x_s}$. Let us represent the ratio of the charged to uncharged coenzyme $\alpha$ as $\Gamma_\alpha = \frac{x_{\alpha'}}{x_\alpha}$, and represent the free energy of reaction $r$, coupled to coenzyme pair $\alpha$ as:

$$\Delta G'_r = \Delta G'^{\dagger}_r + RT\ln\Gamma_\alpha$$

For each reaction $r$, the ratio of the forward flux $v^f_r$, to reverse flux, $v^r_r$ can be represented as a function of the reaction driving force (2), such that:

$$\frac{v^f_r}{v^r_r} = e^{-\Delta G'_r/RT}$$

Substituting our representation of the change in free energy into the previous expression, we have:

$$\frac{v^f_{r\alpha}}{v^r_{r\alpha}} = \frac{K^\dagger_r}{\Gamma_\alpha}$$

where $K^\dagger_r = e^{-\frac{\Delta G^{\dagger}{}'_r}{RT}}$. We now re-cast the equation in terms of the total flux $v^T_r = v^f_r + v^r_r$ and the net flux $v_r = v^f_r - v^r_r$, such that:

$$\frac{v^T_{r\alpha}}{v_{r\alpha}} = \frac{\frac{K^\dagger_r}{\Gamma_\alpha} + 1}{\frac{K^\dagger_r}{\Gamma_\alpha} - 1}$$

Note that reaction $r$ can be driven either by a strong *coenzyme-dependent* force (proportional to $\frac{1}{\Gamma_\alpha}$) or a strong *coenzyme-independent* force (proportional to $K^\dagger_r$). Let us assume that the total flux, on average and over evolutionary time-scales, is linearly proportional to the enzyme concentration, such that $v^T_{r\alpha} = \kappa_r E_{r\alpha}$, where $\kappa_r$ is the sum of the forward and reverse maximal rate constants. Modeling the total flux term as a linear function of enzyme abundance assumes that the enzyme is operating near saturation for both the substrate and product. This assumption is supported by recent studies that suggest that a large portion of NAD(P)-utilizing enzymes operate near saturation, and this property is conserved across phylogenetically distant species (5). Substituting this term for the enzyme abundance into the previous equation, we have the final functional form of enzyme abundance as a function of flux and thermodynamic drive from coenzyme $\alpha$:

$$E_{r\alpha} = \frac{v_{r\alpha}}{\kappa_r}\left(\frac{K^\dagger_r + \Gamma_\alpha}{K^\dagger_r - \Gamma_\alpha}\right)$$

Note that $v_r$ is not restricted to be positive or negative because if $v_{r\alpha} < 0$, then $K^\dagger_r - \Gamma_\alpha < 0$.

In a complex network, there are reactions that use and produce charged coenzymes. To account for these different types of reactions, we can introduce a stoichiometric coefficient for reaction $r$ for each coenzyme $\alpha$, such that if $s_{r\alpha} > 0$, then the reaction proceeds in the following direction:

$$S_r + \alpha \longrightarrow P_r + \alpha'$$

and if $s_{r\alpha} < 0$, then the reaction proceeds in the charged coenzyme-consuming direction:

$$S_r + \alpha' \longrightarrow P_r + \alpha$$

We can now represent the enzyme abundance to include these stoichiometric relationships using the following formula:

$$E_{r\alpha} = \frac{v_{r\alpha}}{\kappa_r}\left(\frac{K_r^{\dagger} + \Gamma_{\alpha}^{s_{r\alpha}}}{K_r^{\dagger} - \Gamma_{\alpha}^{s_{r\alpha}}}\right)$$

We can extend this expression to model a network of reactions utilizing coenzyme $\alpha$. However, the coenyzme has to be balanced (e.g. the total flux into the coenzyme pool has to equal the total flux out of the coenzyme pool). This property can be expressed using the following single constraint:

$$\sum_r s_{r\alpha} v_{r\alpha} = 0 \quad \forall \alpha$$

In our model, we define reaction directionality indpendent of coenzyme choice, forcing the following constraint: $s_{r\alpha} = s_{r\beta} = ...$ for all coenzymes.

We now assume that the total flux through reaction $r$, $v_r$, is a constant and can be satisfied by the action of multiple enzymes that use one of many coenzymes. This requirement can be expressed using the following constraint:

$$\sum_{\alpha} v_{r\alpha} = v_r \quad \forall r$$

**Transhydrogenase and coenzyme exchange.** In extant cells, chemical moieties on redundant coenzyme pairs can be exchanged using transhydrogenases (E.C. 1.6.1.2) for NADH/NADPH exchange or nucleoside-diphosphate kinases (E.C. 2.7.4.6) for ATP/GTP exchange. Let the reaction $e$ catalyze the exchange between two coenzyme pairs, $\alpha$ and $\beta$, such that:

$$\alpha + \beta' \longrightarrow \alpha' + \beta$$

Under standard molar conditions, this reaction is at thermodynamic equilibrium, and thus the only factor that drives net flux is the difference in the concentration ratios for coenzyme $\alpha$ and $\beta$. This, in turn, yields the following expression for the minimum enzyme abundance required to catalyze the exchange reaction:

$$E_e = \frac{v_e}{\kappa_e}\left(\frac{1 + \Gamma_{\alpha}^{s_{e\alpha}}\Gamma_{\beta}^{s_{e\beta}}}{1 - \Gamma_{\alpha}^{s_{e\alpha}}\Gamma_{\beta}^{s_{e\beta}}}\right)$$

For convenience, for multiple coenzymes ordered by a greek index, $\alpha, \beta, \gamma, \ldots$, we include the following constraint that fixes the reaction directionality of each transhydrogenase reaction: $\Gamma_{\alpha} \leq \Gamma_{\beta} \leq \Gamma_{\gamma} \leq \ldots$. This constraint ensures that for the transhydrogenase reaction $e$ that couples coenzyme pair $\alpha$ to $\beta$, $s_{e\alpha} = 1$ and $s_{e\beta} = -1$. This constraint then ensures the reaction flux $v_e \geq 0$, and the enzyme abundance for reaction $e$ becomes:

$$E_e = \frac{v_e}{\kappa_e}\left(\frac{\Gamma_{\beta} + \Gamma_{\alpha}}{\Gamma_{\beta} - \Gamma_{\alpha}}\right)$$

**Enzyme cost minimization.** We wanted to explore whether distributing flux across two coenzyme pools (rather than one) could reduce the protein abundance required to sustain net flux $v_r$ for each reaction $r$ in a network. Including the terms for minimum enzyme abundance required to catalyze reaction $r$ coupled to coenzyme $\alpha$, $E_{r\alpha}$, and each transhydrogenase reaction, $E_e$, we define the following optimization problem to minimize $E_{tot} = \sum_{r\alpha} E_{r\alpha} + \sum_e E_e$, such that:

$$\begin{aligned}
\underset{v_{r\alpha}, v_e, \Gamma_{\alpha}, \Gamma_{\beta}, \ldots}{\text{minimize}} \quad & \sum_{r\alpha} \frac{v_{r\alpha}}{\kappa_r}\left(\frac{K_r^{\dagger} + \Gamma_{\alpha}^{s_{r\alpha}}}{K_r^{\dagger} - \Gamma_{\alpha}^{s_{r\alpha}}}\right) + \sum_e \frac{v_e}{\kappa_e}\left(\frac{\Gamma_{\beta} + \Gamma_{\alpha}}{\Gamma_{\beta} - \Gamma_{\alpha}}\right) \\
\text{subject to} \quad & \sum_{\alpha} v_{r\alpha} = v_r \quad \forall r \\
& \sum_r s_{r\alpha} v_{r\alpha} + \sum_e s_{e\alpha} v_e = 0 \quad \forall \alpha \\
& \Gamma_{\alpha} \leq \Gamma_{\beta} \quad \forall \alpha, \beta
\end{aligned}$$

Note that this is a non-linear optimization problem, with linear equality and inequality constraints. The first set of constraints simply forces the sum of each coenzyme-coupled reaction flux to adhere to a total reaction flux demand, that can be fulfilled by set of coenzymes. The second set of constraints ensures that each coenzyme is at steady state, where $\frac{\partial x_{\alpha}}{\partial t} = 0$. The last set of constraints ensures $v_e \geq 0$, for ordered sets of coenzyme pairs $\{\alpha, \beta\}$.

If we model the optimization assuming a fixed concentration gradient ratios, we can aggregate the kinetic and thermodynamic terms into the following $w_{r\alpha} = \frac{1}{\kappa_r}\left(\frac{K_r^\dagger + \Gamma_\alpha^{s_{r\alpha}}}{K_r^\dagger - \Gamma_\alpha^{s_{r\alpha}}}\right)$, and $w_e = \frac{1}{\kappa_e}\left(\frac{\Gamma_\beta + \Gamma_\alpha}{\Gamma_\beta - \Gamma_\alpha}\right)$ and the optimization problem reduces to a simple linear program:

$$\begin{aligned} \underset{v_{r\alpha}, v_e}{\text{minimize}} \quad & \sum_{r\alpha} w_{r\alpha} v_{r\alpha} + \sum_e w_e v_e \\ \text{subject to} \quad & \sum_{\alpha} v_{r\alpha} = v_r \quad \forall r \\ & \sum_r s_{r\alpha} v_{r\alpha} + \sum_e s_{e\alpha} v_e = 0 \quad \forall \alpha \end{aligned}$$

For models with $N$ reactions and $C$ coenzymes, the total number of free parameters is $NC + C(C-1)/2$ and the total number of constraints is $N + C$.

**Parameter sampling.** We first define parameter regimes one could explore. The first regime is where all enzyme-catalyzed reactions do not rely on a specific coenzyme pair for feasible reaction flux. For the this scenario, the flux through reaction $r$ can be achieved using a single coenzyme or multiple coenzyme pairs. The second regime is where the reaction directionality can be driven by coenzyme $\alpha$. Note that while parameter sampling schemes are discussed for each regime below, we focused our analysis on the first regime because our flux analysis results suggested that a single coenzyme model for *E. coli* core metabolism remained feasible.

For both scenarios, we first sample the stoichiometric coefficients, net fluxes through each reaction $r$, the coenzyme-independent thermodynamic drive, and maximual turnover rates using the following procedure. We first sample the coenzyme stoichiometry, $s_r = s_{r\alpha} = s_{r\beta} = ...$ for each reaction $r$, such that:

$$s_r \sim 2 \times \text{Binomial}(p_s) - 1$$

where $p_s$ sets the proportion of coenzyme-producing to coenzyme consuming reactions. For all simulations presented in Fig. 4 in the main text, and Fig. S5, we set the $p_s = 0.5$. We next define the total flux through the coenzyme pool, $v$, such that the production of all charged coenzymes is balanced by the consumption of all charged coenzymes. Let reaction $p \in \mathcal{R}_p$ denote a reaction the proceeds in the charged coenzyme-producing direction, such that $s_p = s_{p\alpha} = ... = 1$, and reaction $c \in \mathcal{R}_c$ denote a reaction that proceeds in the charged coenzyme consuming direction, such that $s_c = s_{c\alpha} = ... = -1$. We now define the total flux through the coenzyme pools, $v$ using the following relations:

$$v = \sum_{p \in \mathcal{R}_p} v_p = \sum_{c \in \mathcal{R}_c} v_c$$

To obtain proportions of all reaction fluxes, $v_c$ and $v_p$, we use a Dirichlet distribution using the following procedure:

$$v_c, v_p \sim v \times \text{Dirichlet}(a)$$

where $a$ is a concentration parameter of dimensionality $|\mathcal{R}_c|$ and $|\mathcal{R}_p|$ for charged coenzyme consuming and charged coenzyme producing reactions, respectively. For simulations presented in Fig. 4 in the main text, and Fig. S5, we set the total flux to be equal to $v = 100$, and the concentration parameter $a = 1$. We next sample the coenzyme-independent thermodynamic driving force, and the maximal turnover rates using the following log-normal distributions:

$$\log(K_p^\dagger) \sim \mathcal{N}(\mu_{K_p}, \sigma_{K_p}^2)$$

$$\log(K_c^\dagger) \sim \mathcal{N}(\mu_{K_c}, \sigma_{K_c}^2)$$

$$\log(\kappa_r), \log(\kappa_e) \sim \mathcal{N}(\mu_\kappa, \sigma_\kappa^2)$$

***Scenario 1: Reaction directionality is not controlled by coenzyme choice.*** This first scenario is when the thermodynamic gradient provided by the choice of the coenzyme, $\Gamma_\alpha$ does not govern the directionality of the reaction, meaning that $K_r^\dagger$ supplies the free energy required to ensure reaction $r$ goes in the prescribed direction. For this scenario, without loss of generality, we can constrain each reaction to be positive, such that $v_{r\alpha} \geq 0$ for all coenzymes and reactions. By definition, this constrains $\Gamma_\alpha$ such that $K_r^\dagger - \Gamma_\alpha^{s_{r\alpha}} > 0$ for all reactions and coenzymes. For a model with a single coenzyme, we can define the bounds on the feasible ranges of $\Gamma_\alpha$ with the following constraint:

$$\max_{c \in \mathcal{R}_c} \left\{ \frac{1}{K_c^\dagger} \right\} < \Gamma_\alpha < \min_{p \in \mathcal{R}_p} \left\{ K_p^\dagger \right\}$$

However, when multiple coenzyme are available, this constraint only applies to reactions with non-zero flux, such that $v_{r\alpha} \neq 0$. To alleviate this problem, we perform a grid search by varying $\Gamma_\alpha$ for each coenzyme $\alpha$ and determine whether there is a feasible solution before performing linear optimization.

***Scenario 2: Reaction directionality can be controlled by coenzyme choice.*** In the following sampling procedure, reaction directionality can be influenced by coenzyme choice. For this procedure, we define a single distribution of coenzyme-independent driving forces across all reactions, rather than distinguishing between coenzyme-consuming and coenzyme-producing reactions. For each reaction $r$, we first sample the effective equilibrium constant, we sample $\log(K_r^\dagger) \sim \mathcal{N}(\mu_K,\, \sigma_K^2)$, and the catalytic rates $\log(\kappa_r) \sim \mathcal{N}(\mu_\kappa,\, \sigma_\kappa^2)$. We then choose a range of feasible $\Gamma_\alpha$, such that for each coenzyme $\alpha$, there is at-least one reaction in the coenzyme-charging and un-charging direction.

### Minimal enzyme abundance for a simple reaction network

Here we provide an analysis of a simple reaction network, where a single reaction produces the charged coenzyme, $v_p$ and a single reaction flux consumes the charged coenzyme, $v_c$. For this model, the reaction fluxes must be balanced, so $v_c = v_p = v$. The total enzyme required to catalyze reaction flux $v$ is:

$$E = \frac{v}{\kappa_p}\frac{K_p^\dagger + \Gamma}{K_p^\dagger - \Gamma} + \frac{v}{\kappa_c}\frac{K_c^\dagger\Gamma + 1}{K_c^\dagger\Gamma - 1}$$

We can therefore compute the gradient relative to $\Gamma$, such that:

$$\frac{\partial E}{\partial \Gamma} = v\left(\frac{1}{\kappa_p}\frac{2K_p^\dagger}{\left(K_p^\dagger - \Gamma\right)^2} - \frac{1}{\kappa_c}\frac{2K_c^\dagger}{\left(\Gamma K_c^\dagger - 1\right)^2}\right)$$

Setting the derivative to zero, $\frac{\partial E}{\partial \Gamma} = 0$ and solving for $\Gamma$, we obtain the following stationary solution, $\Gamma^*$:

$$\Gamma^* = \frac{K_p^\dagger K_c^\dagger(q_p - q_c) \pm \sqrt{K_p^\dagger K_c^\dagger q_p q_c (K_p^\dagger K_c^\dagger - 1)^2}}{K_c^\dagger(K_p^\dagger K_c^\dagger q_p - q_c)}$$

where $q_p = \frac{v}{\kappa_p}$ and where $q_c = \frac{v}{\kappa_c}$. Note that when the kinetic terms are equivalent for the forward and reverse reactions, such that $q_c = q_p$, this term reduces to single stationary solution:

$$\Gamma^* = \sqrt{\frac{K_p^\dagger}{K_c^\dagger}}$$


### Supplementary Figures

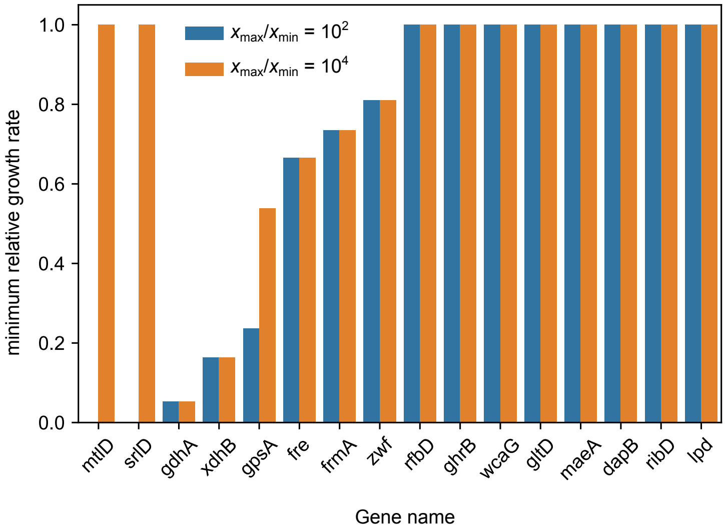


**Fig. S1. Metabolite concentration ranges influence growth rates of coenzyme-swapping mutations in *E. coli*.** We altered the modeling procedure presented in Fig. 2a-b of the main text to reduce the feasible range of metabolite concentrations, such that the maximum metabolite to minimum metabolite concentration ratio ($\frac{x_{\text{max}}}{x_{\text{min}}}$) was reduced from four orders of magnitude to two orders of magnitude. For each coenzyme mutation ($x$-axis), we computed maximum growth rates across all 109,521 media conditions, and plotted the minimum relative growth rate ($\frac{\mu_{\text{mut}}}{\mu_{\text{wt}}}$) across all conditions ($y$-axis).

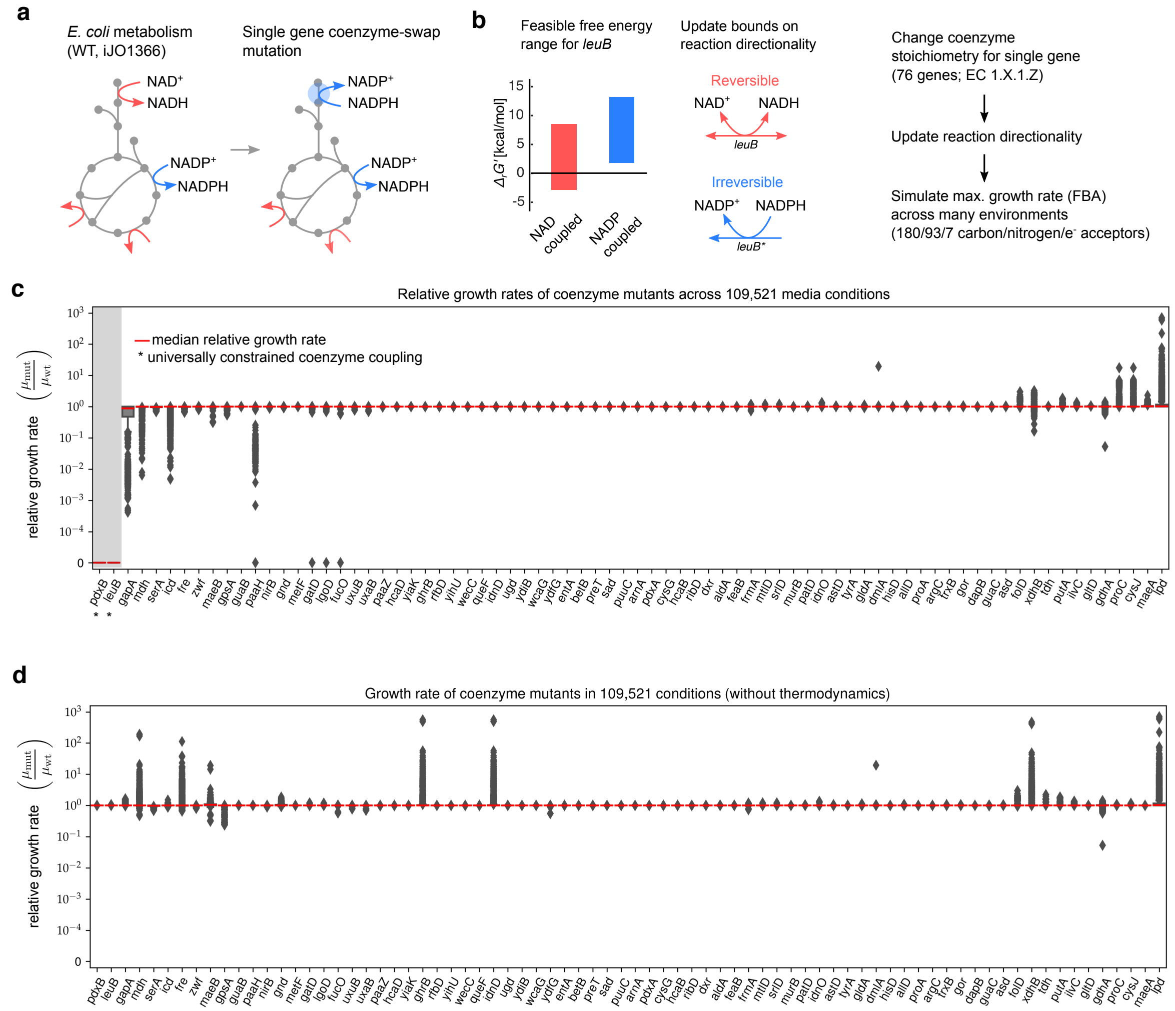


**Fig. S2. Relative growth rates of various coenzyme mutations with and without thermodynamic constraints.** (a) A detailed description of the computational experiments performed in Fig. 2 in the main text. We swapped individual NAD(P)-dependent oxidoreductases with the opposite coenzyme by first altering the stoichiometric dependencies for each individual reaction in the *E. coli* metabolic model, iJO1366. (b) We updated reaction bounds based on computed ranges of reaction free energies (see Methods), where reactions were irreversible if the free energy range did not span both positive and negative values. (c) Each mutant is plotted on the x-axis and the relative growth rate ( $\frac{\mu_{\mathrm{mut}}}{\mu_{\mathrm{wt}}}$ ) , is plotted on the $y$-axis. The grey box denotes the coenzyme swaps where no growth was observed in 100 % media.(d) We repeated the procedure described in (c), but without updating thermodynamic constraints using knowledge of NAD(P)/NAD(P)H ratios. Notably, all essential coenzyme dependencies observed in (c) are no longer observed.

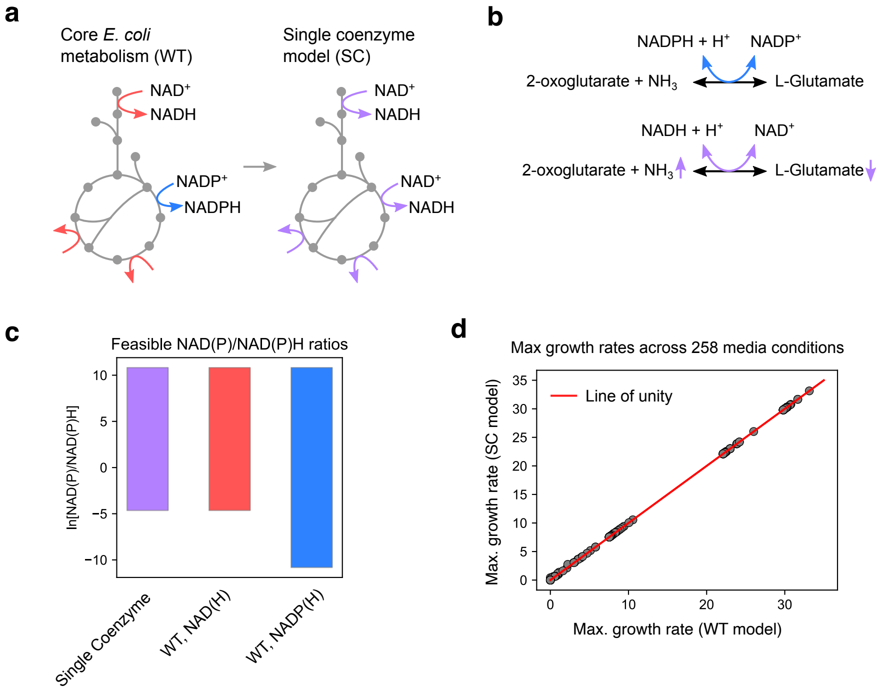


**Fig. S3. Modeling the consequences of replacing all NADP(H) coupled reactions with NAD(H) in *E. coli* core metabolism.** (a) We modified the reduced genome-scale metabolic model irJO1366 by replacing all NADP(H)-coupled reactions with NAD(H), and removed the transhydrogenase reaction and simulated growth in aerobic conditions with glucose as the sole carbon source. (b) Example of a reaction, glutamate dehydrogenase, which is natively originally coupled to NADP(H), that maintained thermodynamic feasibility by changing co-substrate concentrations (e.g., by decreasing L-glutamate and increasing 2-oxoglutarate and ammonia). (c) Feasible ranges of NADP)+/NAD(P)H ratios (y-axis) for the single coenzyme model, and the wildtype model at maximum growth rate. (d) A scatterplot that shows the maximum growth rates of the wild-type (WT, x-axis) vs. single coenzyme model (SC, y-axis) growth rate on a variety of carbon and nitrogen sources, in both anaerobic and aerobic environments.

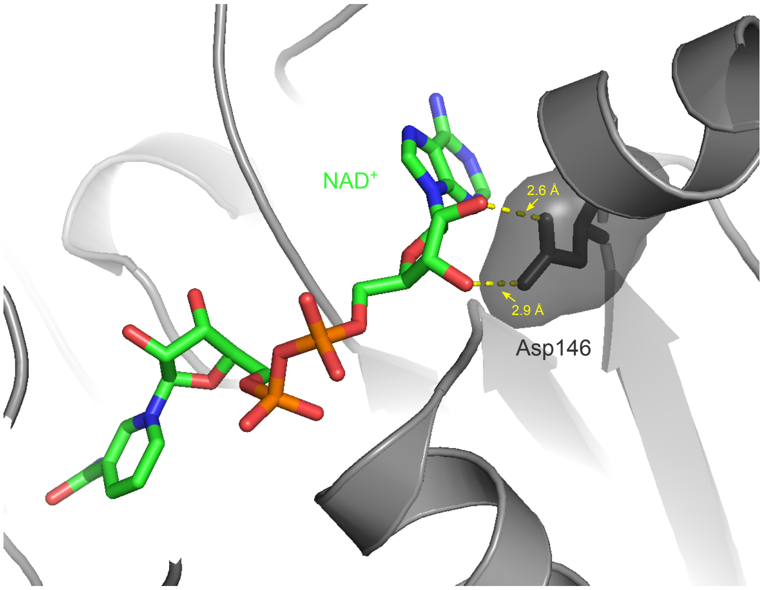


**Fig. S4. NAD-selectivity in D-Erythronate-4-Phosphate Dehydrogenase** The crystal structure of S. enterica D-Erythronate-4-Phosphate Dehydrogenase (encoded by *pdxB*) complexed with NAD+ (PDB 3OET), demonstrating that Asp146 interacts with the 2´ and 3´ hydroxyl groups on the ribose moiety of NAD(H), preventing NADP(H) binding.

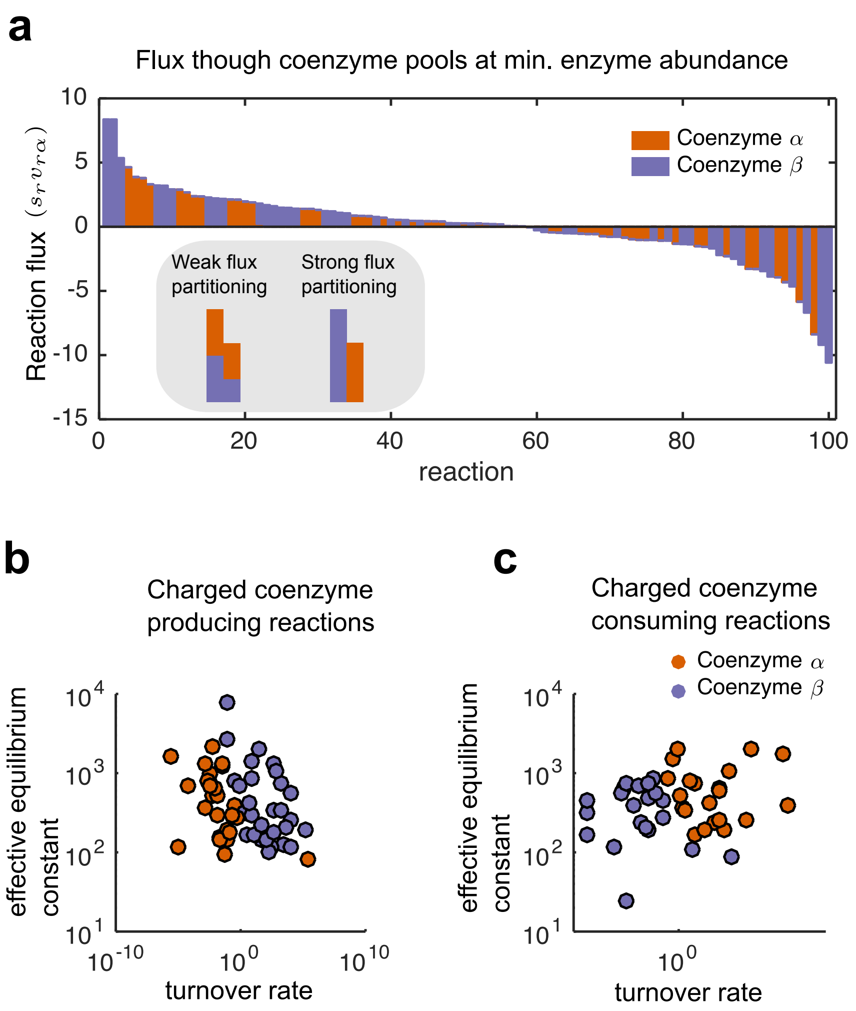


**Fig. S5. Enzyme specificity emerges from proteome expression minimization** a. At the point of minimum total enzyme expression, we plotted the flux (y-axis) for each reaction (x-axis) as a stacked bar plot, where reaction flux through coenzyme $\alpha$ and $\beta$ are shown in orange and purple, respectively. b-c For each reaction, we plotted the sampled turnover rate (x-axis) vs. the sampled effective equilibrium constant (y-axis), and colored each reaction by the preferred coenzyme. For the charged coenzyme producing reactions (b), enzymes with stronger driving forces and faster kinetics were preferentially used coenzyme $\beta$, where these reactions were responsible for operating against a concentration gradient.

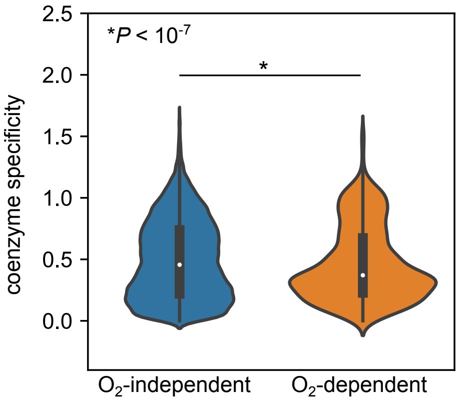


**Fig. S6. Coenzyme specificity is higher in enzymes that catalyze oxygen-independent reactions vs. oxygen-dependent reaction** We computed binding likelihoods for NAD(H) ($L_{r,\text{NAD}}$) and NADP(H) ($L_{r,\text{NADP}}$) for each Rossman folds (denoted by $r$) that were predicted to bind at-least one coenzyme and were not predicted to only bind FAD(H2) ($n = 130, 215$). Binding likelihoods were converted to specificities for each Rossman fold ($s_r$) using the following formula:

$$s_r = |\frac{L_{r,\text{NAD}}}{L_{r,\text{NADP}}}|$$

We then plotted distributions of $s_r$ for oxygen-independent ($n = 126, 718$) and oxygen-independent ($n = 3497$) Rossman folds, and found that folds from oxygen-independent reactions had higher specificity than folds from oxygen-dependent reactions ($P < 10^{-7}$, Mann-Whitney U-Test).
